# Sister chromatid exchanges induced by perturbed replication are formed independently of homologous recombination factors

**DOI:** 10.1101/2021.09.17.460736

**Authors:** Anne Margriet Heijink, Colin Stok, David Porubsky, Eleni M. Manolika, Yannick P. Kok, Marieke Everts, H. Rudolf de Boer, Anastasia Audrey, Elles Wierenga, Victor Guryev, Diana C.J. Spierings, Puck Knipscheer, Arnab Ray Chaudhuri, Peter M. Lansdorp, Marcel A.T.M. van Vugt

**Affiliations:** Department of Medical Oncology, University Medical Center Groningen, University of Groningen, the Netherlands, Hanzeplein 1, 9713GZ, Groningen, the Netherlands; European Institute for the Biology of Ageing, University Medical Center Groningen, University of Groningen, Hanzeplein 1, 9713GZ, Groningen, the Netherlands; Department of Molecular Genetics, Erasmus University Medical Center, 3000 CA Rotterdam, The Netherlands; Oncode Institute, Hubrecht Institute-KNAW and University Medical Center Utrecht, Utrecht, The Netherlands; Terry Fox Laboratory, BC Cancer Agency, Vancouver, BC, V5Z 1L3, Canada; Department of Medical Genetics, University of British Columbia, Vancouver, BC, V6T 1Z4, Canada; Lunenfeld-Tanenbaum Research Institute, Mount Sinai Hospital, 600 University Avenue, Toronto, ON, M5G 1X5, Canada; Department of Molecular Genetics, University of Toronto, 1 King’s College Circle, Toronto, ON, M5S 1A8, Canada; Department of Genome Sciences, University of Washington, Seattle, WA, USA

**Keywords:** PARP inhibitors, synthetic lethality, genome stability, HR, SCE, cross-over, DNA under-replication

## Abstract

Sister chromatid exchanges (SCEs) are products of joint DNA molecule resolution, and are considered to form through homologous recombination (HR). Indeed, upon generation of irradiation-induced DNA breaks, SCE induction was compromised in cells deficient for canonical HR factors BRCA1, BRCA2 and RAD51. Contrarily, replication-blocking agents, including PARP inhibitors, induced SCEs independently of BRCA1, BRCA2 and RAD51. PARP inhibitor-induced SCEs were enriched at common fragile sites (CFSs), and were accompanied by post-replicative single-stranded DNA (ssDNA) gaps. Moreover, PARP inhibitor-induced replication lesions were transmitted into mitosis, suggesting that SCEs originate from mitotic processing of under-replicated DNA. We found that DNA polymerase theta (POLQ) was recruited to mitotic DNA lesions, and loss of POLQ resulted in reduced SCE numbers and severe chromosome fragmentation upon PARP inhibition in HR-deficient cells. Combined, our data show that PARP inhibition generates under-replicated DNA, which is transferred into mitosis and processed into SCEs, independently of canonical HR factors.

## Introduction

Double-stranded DNA breaks (DSBs) are toxic DNA lesions that can lead to cell death or genomic alterations if left unrepaired. Cells have evolved multiple DNA repair mechanisms to deal with DNA breaks (Ciccia and Elledge, 2010; Jackson and Bartek, 2009). In G1 phase of the cell cycle, DNA breaks are predominantly repaired through non-homologous end-joining (NHEJ), which involves ligation of DNA ends independently of sequence homology and frequently results in the introduction of small indels across the break site (Lieber, 2010). In contrast, when DNA has been replicated during S-phase, the sister chromatids can be used as templates for error-free repair through homologous recombination (HR) (Wyman et al., 2004). Cyclin-dependent kinase (CDK)-mediated activation of CtIP promotes the resection of the broken DNA ends by BRCA1/BARD1 and the MRE11/RAD50/NBS1 (MRN) complex, generating single-stranded DNA (ssDNA) overhangs. The resected ends are poor substrates for NHEJ, marking a ‘point of no return’ for HR. Subsequently, BRCA2 promotes loading of RAD51 recombinase onto ssDNA stretches (Sharan et al., 1997; Yang et al., 2002). RAD51 monomers are assembled into nucleoprotein filaments, which ultimately perform the homology search and invasion of the repair template (Petukhova et al., 1998; Sigurdsson et al., 2002). Upon finding homology with the sister chromatid, DNA synthesis takes place via synthesis-dependent strand annealing (SDSA) or through the formation of a joint DNA molecule known as a Holliday junction (HJ). In order to allow faithful chromosome segregation, HJs need to be cleaved before the onset of mitosis (West et al., 2015). HJs can be either ‘dissolved’ by the BLM/RMI1/RMI2/TopIIIa (BTR) complex or ‘resolved’ by the SLX1/SLX4/MUS81/EME1 complex or the GEN1 nuclease (West et al., 2015). Upon completion of DNA synthesis by SDSA and after BTR-mediated dissolution of a HJ, the two ends of the DNA break are rejoined to the original sister chromatid, giving rise to a ‘non-crossover’ event. Alternatively, when the DNA ends of opposing sister chromatids are rejoined, this results in a ‘crossover’ event or ‘sister chromatid exchange (SCE)’ (Castor et al., 2013; Fekairi et al., 2009; Garner et al., 2013; Svendsen et al., 2009; Wyatt et al., 2013). Thus, whereas HJ dissolution exclusively gives rise to non-crossover events (Wu and Hickson, 2003), HJ resolution can give rise to either non-crossover events or crossover end products.

Cells that lack functional BRCA1 or BRCA2, as for instance observed in hereditary breast or ovarian cancers, are defective in homologous recombination and display high levels of genomic instability (Davies et al., 2017; Jonkers et al., 2001; Liu et al., 2007). Due to their DNA repair defect, HR-deficient cancer cells display enhanced sensitivity to DNA damaging agents, including DNA cross-linking agents such as cisplatin (Byrski et al., 2010; Silver et al., 2010). Particularly, HR-deficient cells are sensitive to inhibition of PARP1, an enzyme involved in DNA single strand break repair (Bryant et al., 2005; Farmer et al., 2005). The synthetic lethal interaction between BRCA deficiency and PARP1 inhibitors was initially explained by accumulation of single-strand DNA breaks due to PARP inhibition, which are converted into DSBs that are toxic in the absence of HR repair. However, PARP inhibitors also trap PARP molecules onto DNA (Murai et al., 2012). PARP trapping induces stalling and collapse of replication forks (Schoonen et al., 2017; Zellweger et al., 2015), which are resolved, at least in part, by the HR machinery (Arnaudeau et al., 2001). As a consequence, PARP inactivation leads to elevated levels of SCEs, which are considered products of HR (Ito et al., 2016; Ménissier De Murcia et al., 1997; Wang et al., 1997).

The prevailing model in literature describes a requirement for HR components, including RAD51, in the formation of spontaneous and mutagen-induced SCEs (Sonoda et al., 1999; Wilson and Thompson, 2007). Here we show that, in contrast to irradiation (IR)-induced lesions, DNA lesions induced by replication-blocking agents, including PARP inhibitors, give rise to under-replicated DNA regions in mitosis, which are processed into SCEs independently of canonical HR.

## Results

### Olaparib induces sister-chromatid exchanges in BRCA2-proficient and deficient cancer cells

To study the effects of PARP inhibition on the induction of sister-chromatid exchanges (SCEs), we employed Strand-seq (Falconer et al., 2012; Sanders et al., 2017), which allows single cell sequencing of the DNA template strand and genome-wide mapping of SCEs (van Wietmarschen et al., 2018). Murine tumor-derived *Tp53*^−/−^*Brca2*^−/−^ KB2P3.4 cells and *Brca2*-reconstituted *Tp53*^−/−^*Brca2*^IBAC^ KB2P3.4R3 cells (Evers et al., 2010) were treated with the PARP inhibitor olaparib, and libraries were prepared for Strand-seq analysis (Fig. 1A). A significant increase in the number of SCEs was observed in *Brca2*-proficient KB2P3.4R3 cells in response to PARP inhibitor treatment (Fig. 1A, B), in line with previous findings (Ito et al., 2016; Oikawa et al., 1980). Surprisingly however, Strand-seq analysis revealed that olaparib treatment induced a comparable number of SCEs in *Brca2*-deficient KB2P3.4 cells (Fig. 1A, B). Of note, a significant increase in copy number variations (CNVs) was observed in *Brca2*-deficient KB2P3.4 cells (Fig. 1C, Supplementary Fig. S1A), a phenotype consistent with HR-deficiency (Davies et al., 2017). Using differential “harlequin” chromatid staining of metaphase spreads, olaparib-induced SCEs were again observed in the absence of *Brca2*, validating our previous observation (Fig. 1D, E). Moreover, background levels of SCEs were similar in *Brca2*-proficient and *Brca2*-deficient cells, indicating that spontaneous SCEs, like olaparib-induced SCEs, arise independently of *Brca2* (Fig. 1B, E). Thus, both spontaneous and PARP inhibitor-induced SCEs arise in *Brca2*-deficient cells, suggesting that *Brca2* is not essential for SCE formation.

**Figure 1:**
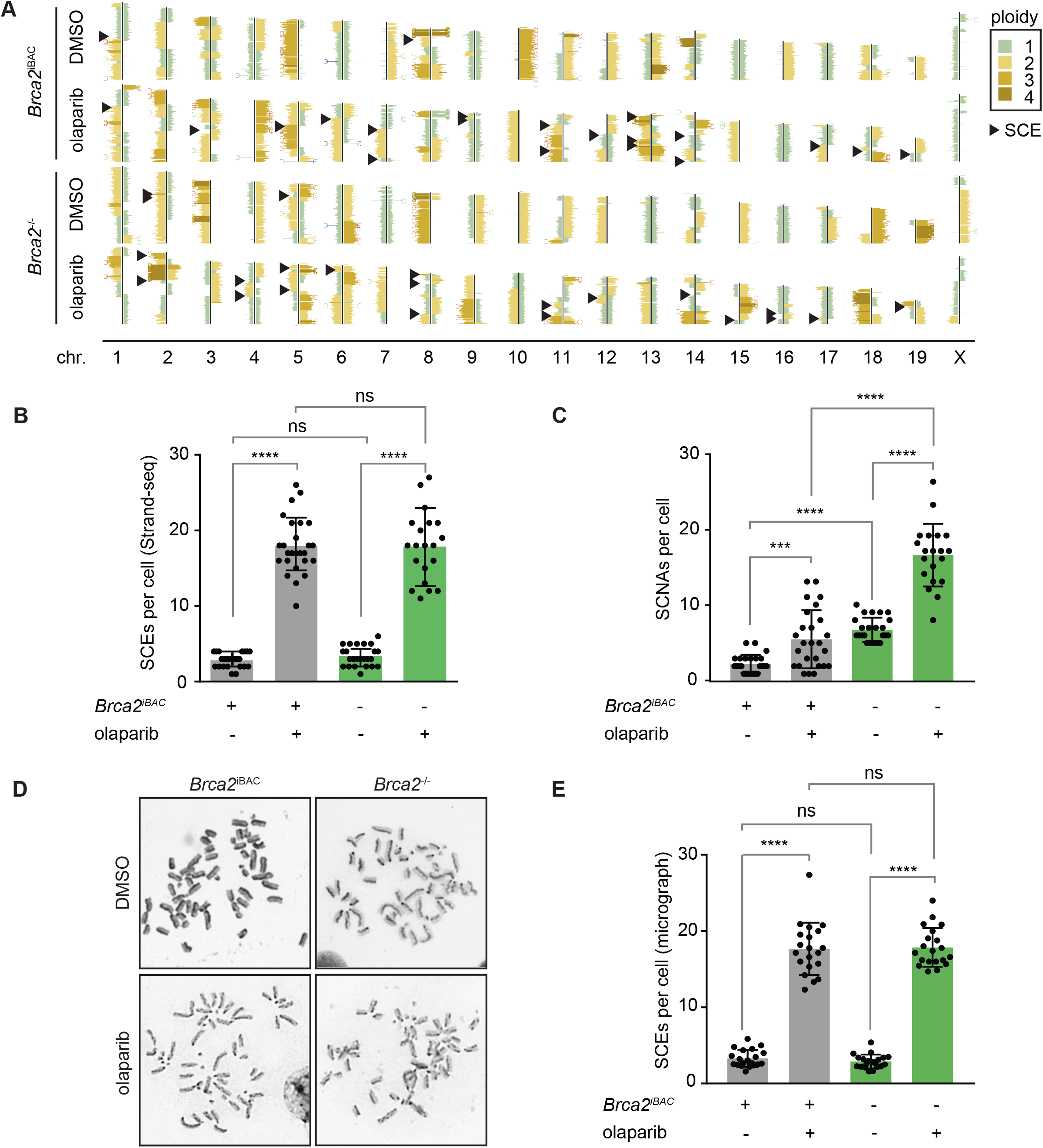
Olaparib-treatment induces sister-chromatid exchanges in *Brca2* wt and *Brca2*-mutant cancer cells. **(A)** Representative Strand-seq libraries of *Brca2*^−/−^ cells or *Brca2*^iBAC^ cells, treated with DMSO (top) or olaparib (bottom). Black arrowheads indicate SCEs. **(B)** *Brca2*^−/−^ cells or *Brca2^iBAC^* cells were treated with DMSO or olaparib. Quantification of SCEs per cell was determined in n#23 (*Brca2*^*iBAC*^, DMSO), n=26 (*Brca2*^*iBAC*^, olaparib), n#24 (*Brca2*^−/−^, DMSO) and n#19 (*Brca2*^−/−^, olaparib) libraries. Error bars represent standard deviation. **(C)** *Brca2*^−/−^ cells or *Brca2*^*iBAC*^ cells were treated with DMSO or olaparib as for panel B. CNVs per cell were quantified. Means and standard deviations are presented. **(D, E)** *Brca2*^−/−^ cells or *Brca2*^*iBAC*^ cells were treated with DMSO or olaparib, and SCEs were quantified by microscopy analysis of at least 30 metaphase spreads per condition (panel D). Averages and standard deviation are presented (panel E). Statistics in panels (B, C, E) were performed using unpaired two-tailed t-tests (ns: p > 0.05, *: p < 0.05, **: p < 0.01, ***: p < 0.001, ****: p < 0.0001).

### Olaparib-induced SCEs arise independently of canonical HR factors in a dose-dependent manner

To investigate whether BRCA2-independent SCEs can be observed in other cell lines, we introduced shRNAs targeting BRCA2 in untransformed human *TP53*^−/−^ RPE-1 cells (Fig. 2A), and used sublethal concentrations of olaparib in these cells (Fig. S2A). We observed a significant and dose-dependent increase in the number of SCEs upon treatment with olaparib (Fig. 2B, S2B), which again occurred independently of BRCA2 (Fig. 2B). Interestingly, induction of SCEs by γ-irradiation (IR) was completely dependent on BRCA2 (Fig. 2C), in line with previous reports (Conrad et al., 2011). To test whether olaparib-induced SCEs also arise independently of other canonical HR components, we depleted BRCA1, an upstream HR regulator, and RAD51, the main recombinase responsible for strand invasion (Fig. 2A). In line with our results in BRCA2-depleted cells, olaparib-induced SCEs were also observed in cells expressing shRNAs against BRCA1 or RAD51 (Fig. 2B). In stark contrast, IR-induced SCEs were fully dependent on BRCA1 and RAD51 (Fig. 2C), indicating that the processing of IR-induced and olaparib-induced lesions show different dependencies on canonical HR factors. Furthermore, proteins involved in the resolution of joint molecules downstream in the HR pathway, including MUS81, ERCC1 and SLX1-SLX4 were also not required for the formation of olaparib-induced SCEs in RPE-1 cells, whereas these proteins were required for IR-induced SCEs (Fig. S2C-E).

**Figure 2:**
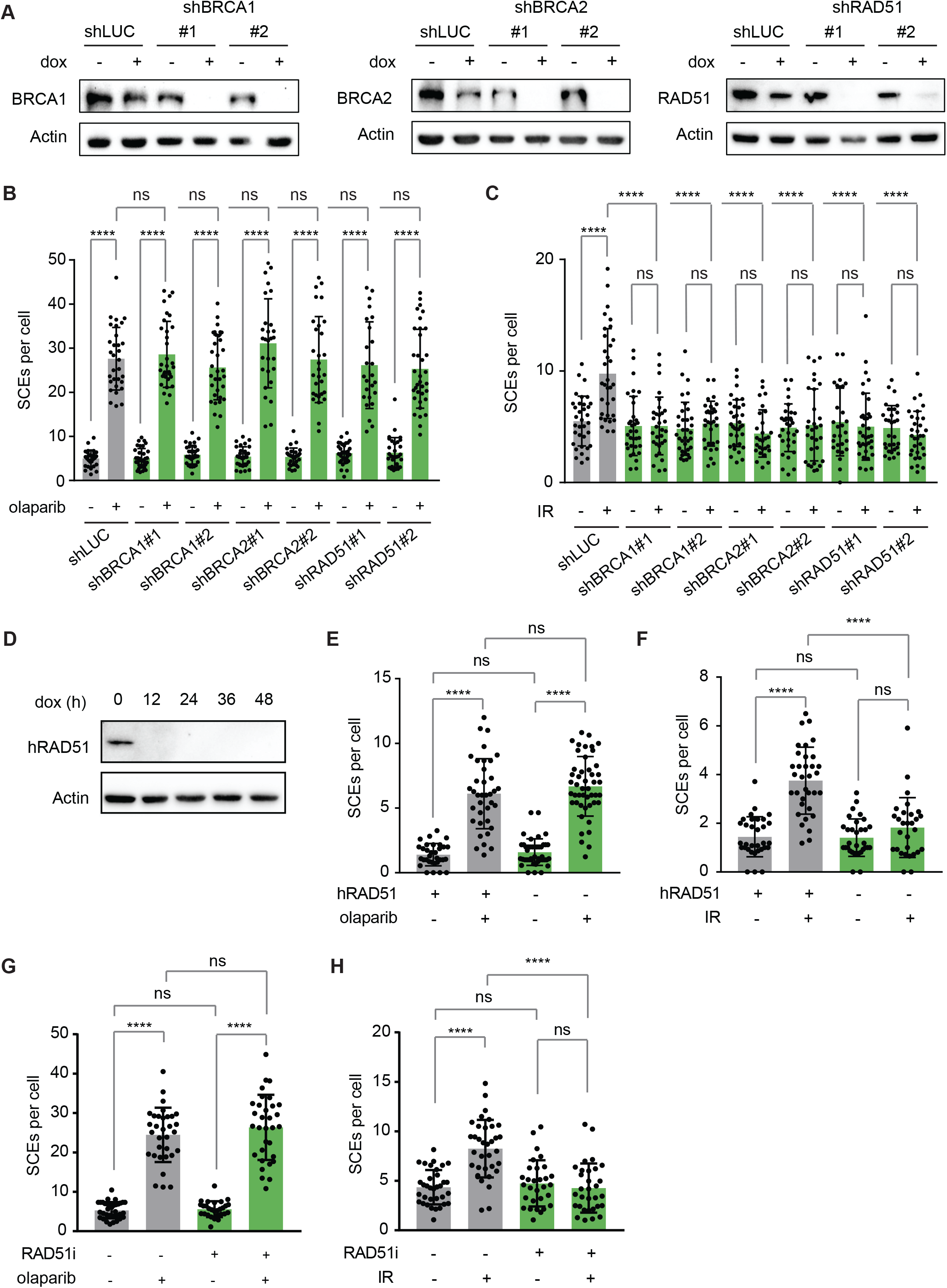
HR-independent induction of SCEs upon PARP inhibitor treatment. **(A)** RPE-1-*TP53*^−/−^ cells with indicated dox-inducible shRNAs were treated with doxycycline for 48 h and immunoblotted for indicated proteins. **(B, C)** RPE-1-*TP53*^−/−^ cells with indicated dox-inducible shRNAs were treated with doxycycline for 48 h, and subsequently treated with olaparib for 48 hours (panel B) or 2 Gy irradiation (panel C). SCEs were quantified by microscopy analysis of at least 30 metaphase spreads per condition. Means and standard deviations are plotted. **(D-F)** DT40 *RAD51*^−/−^ cells harboring a dox-repressed hRad51 transgene were treated with doxycycline for indicated time periods, and immunoblotted for Rad51 and Actin (panel D) or treated with olaparib (panel E) or irradiation (panel F). SCEs were quantified by microscopy analysis of at least 30 metaphase spreads per condition. **(G, H)** RPE1-*TP53*^−/−^ cells were incubated with olaparib (panel G) or IR (panel H) in the absence or presence of the RAD51 inhibitor BO2. SCEs were quantified by microscopy analysis of at least 30 metaphase spreads per condition. Means and standard deviation are plotted. Statistics in panels (B, C, E, F, G, H) were performed using unpaired two-tailed t-tests (ns: p > 0.05, *: p < 0.05, **: p < 0.01, ***: p < 0.001, ****: p < 0.0001).

Since RAD51 has a central role in recombination, SCE formation was assessed in a *RAD51*^−/−^ DT40 chicken B-cell lymphoma cell line as well, which depends on the expression of a doxycycline-repressible hRAD51 transgene for viability (Sonoda et al., 1998). Treatment with doxycycline resulted in robust repression of hRAD51 expression, allowing us to study RAD51-dependent events in these cells (Fig. 2D). Metaphase spreads revealed high numbers of gaps and breaks in these cells, characteristic of RAD51 deficiency (Fig. S2F). Notably, and in accordance with our findings in RPE-1 cells, olaparib treatment in these cells induced formation of SCEs independently of RAD51 (Fig. 2E), whereas IR-induced SCEs were fully dependent on RAD51 (Fig. 2F). In parallel, we used the small molecule inhibitor B02 to inhibit the DNA strand exchange activity of RAD51 (Huang et al., 2011), and again observed SCE induction upon olaparib treatment independently of RAD51 (Fig. 2G). By contrast, RAD51 inhibition completely prevented SCE induction in IR-treated cells (Fig. 2H). Overall, these data show that olaparib-induced SCEs are independent of canonical HR components in multiple HR-deficient cell line models.

### HR-independent SCEs are associated with DNA under-replication

In addition to inhibiting the enzymatic activity of the PARP1/2 enzymes, most PARP inhibitors, including olaparib, are capable of trapping PARP to the DNA (El-Khamisy et al., 2003; Murai et al., 2012). In order to distinguish between the effects of PARP inhibition and PARP trapping, siRNAs against PARP1 were introduced in RPE-1 cells (Fig. 3A). siRNA-mediated depletion of PARP1 induced significantly less SCEs when compared to PARP inhibition (Fig. 3B), suggesting that PARP trapping is the predominant source for the generation of HR-independent SCEs. A range of PARP inhibitors has been developed, with ranging capacity to trap PARP onto DNA (Murai et al., 2012). To further test whether PARP trapping capacity is related to the induction of SCEs, we treated RPE-1 cells with IC25 concentrations of a panel of PARP inhibitors with trapping capacity: olaparib, veliparib and talazoparib (Fig. 3B and S2A). When used at IC25 dose, all three PARP inhibitors induced comparable amounts of SCEs, which were again generated independently of BRCA2 (Fig. 3B). Since PARP-trapping inhibitors have been reported to disturb normal replication fork progression (Bryant et al., 2009; Maya-Mendoza et al., 2018; Schoonen et al., 2017), we investigated whether SCEs also arise upon treatment with other replication-perturbing agents, to exclude effects of the roles of PARP1/2 in DNA repair. A panel of commonly used chemotherapeutics that target DNA replication was tested, including the DNA crosslinking agents mitomycin C (MMC) and cisplatin, and the topoisomerase inhibitors camptothecin (CPT) and etoposide. Although these agents induced SCEs to various extents, very similar amounts of SCEs were observed in control and BRCA2-depleted cells (Fig. 3B). Only for etoposide, a small but significant decrease in SCEs was observed in BRCA2-depleted cells (Fig. 3B). Overall, our data suggests that the capacity to block DNA replication forks is a key determinant in the formation of HR-independent SCEs.

**Figure 3:**
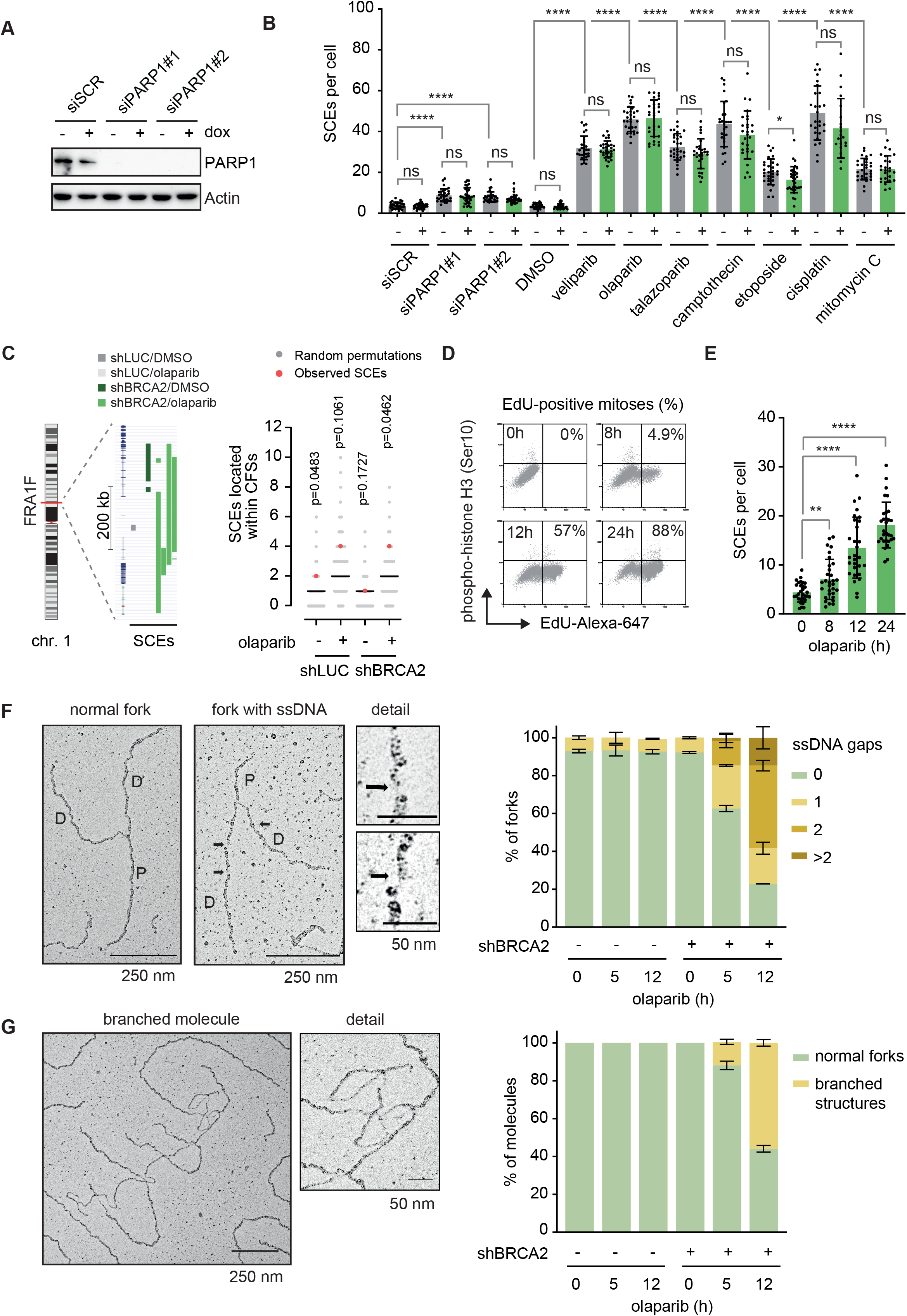
HR-independent SCEs are associated with defective replication. **(A)** RPE-1-*TP53*^−/−^ shBRCA2 cells were pretreated with doxycycline (dox) and transfected with siPARP or control siRNAs. Lysates were immunoblotted for PARP1 and Actin. (B) RPE-1-*TP53*^−/−^ shBRCA2 cells were pre-treated with doxycycline (dox) and subsequently treated with indicated agents for 48 h or transfected with PARP1 siRNAs 48 h before harvest. SCEs were quantified by microscopy analysis of at least 30 metaphase spreads per condition. Averages and standard deviation of 30 mitoses per condition are indicated. **(C)** KBM-7 cells harboring doxycycline-inducible control or BRCA2 shRNAs were pre-treated with doxycycline and subsequently treated with olaparib where indicated. SCEs were mapped using StrandSeq of 64 (shLUC/DMSO), 31 (shLUC/OLA), 50 (shBRCA2/DMSO) and 52 (shBRCA2/OLA) libraries per condition, respectively. Observed SCEs were mapped to CFSs, and compared to 10,000 random permutations. P values indicate deviation of the observed number of SCEs compared to the mean of all permutations. SCE mapping to the common fragile site FRA1F is presented as an illustrative example. **(D)** Doxycycline pre-treated RPE-1-*TP53*^−/−^ shBRCA2 cells were treated with ethynyl deoxyuridine (EdU) for the indicated time periods, and subsequently analyzed for mitotic cells by flow cytometry of phospho-histone H3 (Ser10). Percentages of mitotic cells that were EdU-positive are indicated. **(E)** Doxycycline pre-treated RPE-1-*TP53*^−/−^ shBRCA2 cells were treated with olaparib for the indicated time points, and SCEs were quantified by microscopy analysis of at least 30 metaphase spreads per condition. **(F)** RPE1-*TP53*^−/−^ shBRCA2 cells were pretreated for 48 h with doxycycline (dox) and treated with olaparib for indicated time periods. Representative electron microscopy images are indicated of normal replication forks, and replication forks with ssDNA gaps. Quantification of ssDNA gaps is presented in the right panel. Averages and standard deviations of 60 replication fork per condition are shown. **(G)** RPE1-*TP53*^−/−^ shBRCA2 cells were treated as for panel F. A representative image of branched DNA molecule is shown. Quantification of branched DNA molecules is indicated in the right panel. Averages and standard deviations of 60 replication forks per condition are shown. Statistics in panels (B, E) were performed using unpaired two-tailed t-tests (ns: p > 0.05, *: p < 0.05, **: p < 0.01, ***: p < 0.001, ****: p < 0.0001).

To investigate whether PARP inhibitor-induced SCEs are associated with specific genomic features, we performed Strand-seq in olaparib-treated KBM-7 cells, expressing control or BRCA2-targeting shRNAs. The near haploid karyotype of KBM-7 cells allows for robust mapping of SCEs. In line with our previous observations, PARP inhibition induced SCEs in KBM-7 cells independently of BRCA2 (Fig. S3A). KBM-7 cells displayed deletions, amplifications and copy number variations in BRCA2-depleted cells, which were further increased upon PARP inhibition, underscoring a functional HR defect in these cells (Fig. S3B-D). SCE locations were mapped using HapSCElocatoR, as described previously (Fig. S3E) (Claussin et al., 2017). Genomic locations of SCEs were mapped against the locations of previously described human CFSs (Kumar et al., 2019). An enrichment of olaparib-induced SCEs was observed within CFS regions in BRCA2-deficient cells (Fig. 3C), in line with SCEs being regarded as difficult-to-replicate loci. As an example, in BRCA2-deficient cells treated with olaparib, a substantial number of SCEs were detected within the CFS FRA1B (Fig. 3C). Subsequently, SCEs were mapped against centromeres and telomeres to test enrichment at other difficult-to-replicate regions (Fig. S3F, G). Whereas no SCEs were observed at telomeric regions, olaparib also significantly induced centromeric SCEs in both BRCA2-deficient and control cells (Fig. S3F, G). Moreover, BRCA2-depleted cells showed a significant depletion of SCEs within gene bodies (Fig. S3H). Finally, no significant enrichments were observed at putative G4 structures (Fig. S3I). The observation that SCEs in BRCA2-deficient cells are enriched at CFSs and centromeric regions is in accordance with our hypothesis that these SCEs are associated with DNA replication fork stalling.

To further investigate the relation between DNA replication and SCE formation, RPE-1 cells were treated with olaparib for different time periods, treating cells either during or after S-phase (Fig. 3D, E). EdU incorporation was used to assess the time required for RPE-1 cells to progress from S-phase to mitosis (Fig. 3D). After 8h of EdU incorporation, only 4.9% of all mitoses (phospho-histone H3-positive cells) had incorporated EdU, suggesting that the majority of mitotic cells did not progress through S-phase at the time of collection (Fig. 3D). Accordingly, 8h olaparib treatment induced only a minor number of SCEs (Fig. 3E). In contrast, 12 or 24h EdU treatment resulted in a considerable population of EdU-positive mitotic cells (Fig. 3D), which coincided with significantly larger numbers of SCEs being induced (Fig. 3E). Overall, these data suggest that olaparib needs to be present during S-phase in order to induce SCEs in the following mitosis.

To further assess the effects of PARP inhibition of DNA replication, we used electron microscopy to analyze replication forks (Fig. 3F). Upon olaparib treatment, we observed a significant increase in the amounts of gaps, as judged by the single-stranded DNA patches in the newly synthetized DNA strand (Fig. 3F). In addition, EM analysis revealed complex ‘branched’ replication structures, predominantly in BRCA2-depleted cells upon olaparib treatment (Fig. 3G). Combined, these findings illustrate that PARP inhibition in HR-deficient cells leads to extensive replication perturbation, and that HR-independent SCEs are observed in a range of conditions, with perturbed replication as a shared mechanism-of-action.

### Processing of olaparib-induced DNA lesions during mitosis

We previously reported that olaparib-induced replication lesions are transmitted into mitosis (Schoonen et al., 2017, 2019). To further test whether olaparib treatment induces mitotic DNA lesions in BRCA2-deficient RPE-1 *TP53*^−/−^ cells, we measured the DNA damage markers γH2AX and FANCD2 in mitotic cells (Fig. 4A). Both endogenous and olaparib-induced γH2AX and FANCD2 foci were enriched in BRCA2-depleted cells (Fig. 4A). Since mitotic FANCD2 foci reflect the presence of unresolved under-replicated DNA, we next assessed if olaparib treatment leads to increased mitotic DNA synthesis (MiDAS). We observed an increase in mitotic EdU foci in response to olaparib treatment in BRCA2-depleted cells (Fig. 4B), suggesting that DNA replication in these cells is incomplete at the moment of mitotic entry. In order to further investigate the link between mitosis and SCE formation, RPE-1 cells were treated with an inhibitor of the ATR checkpoint kinase to force cells into mitosis with under-replicated DNA (Fig. 4C). Cells treated with ATR inhibitor showed elevated numbers of SCEs, independently of BRCA2 (Fig. 4E), suggesting that mitotic processing of under-replicated DNA may be the source for these SCEs.

**Figure 4:**
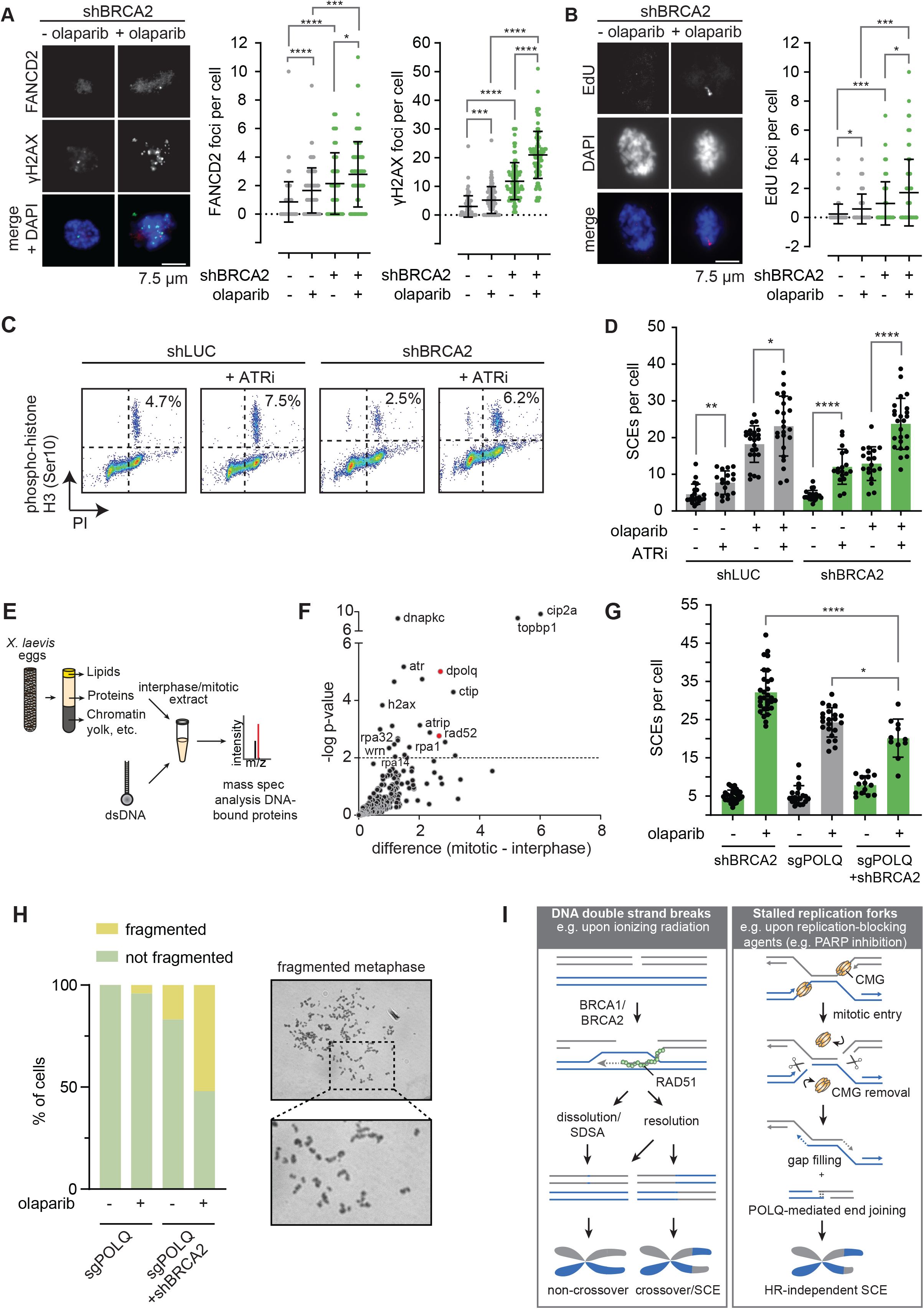
SCEs originate from mitotic processing of under-replicated DNA. **(A)** RPE1-*TP53*^−/−^ shBRCA2 were pretreated with doxycycline (dox), synchronized using RO-3306 for 4 h, and subsequently treated with olaparib where indicated. γH2AX and FANCD2 foci in mitotic cells were quantified by immunofluorescence microscopy. Means and standard deviation of pooled data from three independent experiments are shown, with n#30 mitoses per experiment. **(B)** RPE1-*TP53*^−/−^ shBRCA2 were treated as for panel A. 24 h after olaparib treatment, cells were incubated for 25 min with EdU. Mitotic EdU foci were quantified in n#30 mitoses per experiment. Means and standard deviation of pooled data from three independent experiments are shown. **(D, E)** RPE1-*TP53*^−/−^ shBRCA2 cells were treated with doxycycline (dox) for 48 h, with olaparib for 24 h, with or without the ATR inhibitor VE-821 (ATRi) for 3 h. Cells were treated with colcemid for 3 h before harvesting, fixed and stained for the mitotic marker phospho-Histone-H3 (panel D). In parallel, SCEs were quantified by microscopy analysis of at least 20 metaphase spreads per condition (panel E). **(F, G)** Interphase or mitotic *Xenopus* egg extracts were prepared and incubated with biotin-conjugated blunt-ended DNA oligos (panel F). Proteins associated with DNA oligos were identified by mass spectrometry (panel G). **(H)** RPE1-*TP53*^−/−^ sgPOLQ shBRCA2 were pretreated with doxycycline (dox) and subsequently treated with olaparib where indicated. SCEs were quantified by microscopy analysis of 11-30 mitoses per condition, means and standard deviations are indicated. **(I)** Cells were treated as for panel H, and fragmented DNA was analyzed for 50 mitoses per condition. **(K)** Model of HR-independent SCE formation. Left panel indicates HR-dependent SCE formation after DSB repair. Right panel indicates HR-independent SCE formation by mitotic processing of under-replicated DNA. Statistics in panels (A, B) were performed using Mann-Whitney test, statistics in panel (E, H) were performed using unpaired two-tailed t-tests (ns: p > 0.05, *: p < 0.05, **: p < 0.01, ***: p < 0.001, ****: p < 0.0001).

At the onset of mitosis, stalled replication forks may be cleaved at both leading or both lagging strands, introducing DSBs surrounding the under-replicated genomic region. Since canonical NHEJ is inactivated during mitosis (Benada et al., 2015; Orthwein et al., 2014), we searched for proteins that act on DNA breaks during mitosis. To this end, we used biotin-tagged synthetic DNA structures that resemble DNA double strand breaks, and pulled out associated proteins from interphase or mitotic *Xenopus laevis* egg extracts (Fig. 4E). Mass spectrometry analysis revealed a range of proteins that was enriched on synthetic DNA breaks in mitotic extracts, including Cip2a and Topbp1 (Fig. 4F), which together with Mdc1 are involved in mitotic tethering of mitotic DNA breaks (Adam et al., 2021; Leimbacher et al., 2019). Intriguingly, we also identified the single-strand annealing (SSA) factor Rad52 and the alternative end-joining (alt-EJ) factor Polq (Fig. 4F). Inactivation of RAD52 in RPE-1 cells using CRISPR/Cas9, either alone or in combination with BRCA2 depletion, did not decrease SCEs amounts in olaparib-treated cells (Fig. S4A, B), suggesting that SSA through RAD52 is not responsible for PARP inhibitor-induced SCEs. In contrast, upon CRISPR/Cas9-mediated inactivation of POLQ (Fig. S4C), a decrease in SCEs was observed in BRCA2-deficient RPE-1 cells (Fig. 4G). Moreover, we observed severe fragmentation of mitotic chromosomes, consistent with an inability to process mitotic breaks resulting from under-replicated DNA (Fig. 4H).

Our data fit a model in which transmission of under-replicated DNA into mitosis results in HR-independent SCE formation (Fig. 4I). We hypothesize that upon removal of the replisome at the onset of mitosis, DNA breaks are induced, flanking the under-replicated DNA, consistent with previously reported data (Deng et al., 2019). Subsequently, the ssDNA gaps flanking the under-replicated region are filled, and ligation of the two broken sister chromatids is promoted by POLQ (Fig. 4I). As a consequence of mitotic processing of under-replicated DNA according to this model, an HR-independent SCE is formed, which is predicted to be accompanied by allelic deletions with a size reflecting the extent of under-replication (Fig. 4I). Loss of POLQ may prevent re-ligation of the broken DNA ends, resulting in metaphase chromosome fragmentation (Fig. 4H). This model also predicts that premature mitotic entry with under-replicated DNA is sufficient to induce SCEs, without the need for HR components. Combined, our data suggest that HR-independent SCEs originate from mitotic processing of under-replicated DNA, and suggests the involvement of the alternative end-joining polymerase POLQ.

## Discussion

We here show that agents that perturb DNA replication, including PARP inhibitors, induce sister chromatids exchanges in the absence of canonical HR factors BRCA1, BRCA2 and RAD51. Conversely, these HR components are required for induction of SCEs upon irradiation. Our findings challenge the current dogma that SCEs solely arise as a result of homologous recombination (Dronkert et al., 2000; Polato et al., 2014; Smiraldo et al., 2005; Sonoda et al., 1999; Wilson and Thompson, 2007). Interestingly, models in which formation of SCEs upon replication fork stalling occur independently of HR have been proposed in the past (Ishii and Bender, 1980), although these have lost support in favor of the HR-dependent models over the years. The observation that SCEs form independently of RAD51 is of particular interest. SCEs are considered to involve the formation of joint molecules, which typically requires strand invasion by RAD51. Previously observed replication stress-associated SCEs in *BRCA2* mutant cells were explained by BRCA2-independent RAD51 recruitment (Ray Chaudhuri et al., 2016). Yet, spontaneous and replication-induced SCEs have been reported frequently in cells lacking canonical HR factors (Lambert and Lopez, 2001; Sonoda et al., 1999; Takata et al., 2001; Tutt et al., 2001), underscoring the notion that SCEs can arise independently of HR, and suggesting that PARP-inhibitor induced SCEs and spontaneous SCEs may share common mechanisms. Surprisingly, loss of the RAD51 paralogs RAD54, RAD51C, RAD51D, XRCC2, and XRCC3 has previously been reported to reduce MMC-induced and spontaneous SCEs, although these effects were limited and were attributed to defective Rad51 functioning (Sonoda et al., 1999; Takata et al., 2001).

In good agreement with our data, coordinated cleavage of stalled replication forks in mitosis was recently hypothesized to yield SCEs, at the cost of local deletions (Wu et al., 2021). This process requires that the stalled replication forks flanking the under-replicated DNA are cleaved either at both leading strands, or at both lagging strands (Fig. 4I). Coordinated cleavage of replication forks could be proceeded by TRAIP/p97-dependent unloading of the CMG helicase (Deng et al., 2019), leaving stretches of vulnerable ssDNA at the leading strands. Although the responsible nuclease in processing under-replicated DNA upon PARP inhibition remains elusive, MUS81 has been shown to be active during mitosis (Calzetta et al., 2020; García-Luis and Machín, 2014; Wild et al., 2019; Wyatt et al., 2013), to localize to under-replicated DNA in mitosis (Di Marco et al., 2017), and to act on stalled replication forks in mitosis (Fugger et al., 2021; Zimmermann et al., 2013). However, our data showed that depletion of MUS81 alone was not sufficient to reduce SCEs, possibly due to redundancy with other endonucleases that are active during mitosis (Chan and West, 2014; Wyatt et al., 2013).

Cleavage and end-joining of cleaved under-replicated DNA regions during mitosis would yield SCEs as well as allelic deletions of under-replicated genomic regions. Such allelic deletions have been reported previously at common-fragile sites (Durkin and Glover, 2007). Moreover, the observed submicroscopic deletions that span CFS regions were shown to be flanked by microhomology regions, suggesting the involvement of POLQ-mediated end joining, consistent with the mutational signatures observed in BRCA1/2-deficient tumors (Glover et al., 2017; Stok et al., 2021). Moreover, a role for POLQ in the mitotic processing of stalled replication forks in BRCA-deficient cells fits well with the synthetic lethality between BRCA2 and POLQ, as previously described (Ceccaldi et al., 2015; Mateos-Gomez et al., 2015; Mengwasser et al., 2019).

Interestingly, our observation that mitotic SCEs arise independently of RAD52, rules out that we are looking at a direct end product of RAD52-dependent MiDAS (Bhowmick et al., 2016; Minocherhomji et al., 2015). Interestingly, MiDAS was also described to involve nuclease activity, and this pathway could therefore potentially compete with POLQ-dependent mitotic SCE formation. Besides being dependent on RAD52, the genetic requirement of MiDAS resemble those of break-induced replication (BIR), which involves the formation of joint molecules (Minocherhomji et al., 2015). How MiDAS-mediated joint molecules can be formed during mitosis, when multiple nucleases are active to resolve such structures remains unclear. Interestingly, MiDAS was recently suggested to reflect completion of DNA replication at the end of G2 phase, rather than during mitosis (Mocanu et al., 2021), suggesting that MiDAS and mitotic SCE formation could be consecutive processes.

Although the synthetic lethal interaction between HR loss and PARP inhibition has been validated in various models and has been successfully exploited in the clinic, the fate of PARP inhibitor-induced DNA lesions in HR-deficient cells remains unclear (Helleday, 2011). Although we find that PARP inhibitor-induced DNA lesions are processed into SCEs in HR-deficient cells, we predict that this goes along with accumulation of large deletions and translocations due to mitotic POLQ activity, which may underlie loss of viability observed in these cells. In this context, a requirement for mitotic replisome unloading by TRAIP/p97, fits well with observed sensitization of HR-deficient cells for PARP inhibition upon inactivation of TRAIP (Fugger et al., 2021), and the observation that progression through mitosis promotes PARP inhibitor-mediated cell death (Schoonen et al., 2017). Our observation that PARP inhibitor-induced DNA lesions in HR-deficient cells are transmitted into mitosis also aligns well the recent identification of the DNA tethering factor CIP2A being essential in HR-deficient cells (Adam et al., 2021), and was identified in our proteomics analysis of mitotic factors that bind DNA ends. Further research is warranted to investigate whether tethering of DNA ends is required to ligate DNA ends upon cleavage of stalled replication forks during mitosis.

## Methods

### Cell lines

hTERT-immortalized human retina epithelial RPE-1 cells and HEK293T cells were obtained from ATCC, and were maintained in Dulbecco’s Minimum Essential Media (DMEM, Thermofisher), supplemented with 10% fetal calf serum (FCS, Lonza), 50 units/mL penicillin and 50 μg/mL streptomycin (P/S, Gibco) at 37°C, 20% O_2_ and 5% CO_2_. KBM-7 cells were a kind gift from Thijn Brummelkamp (The Netherlands Cancer Institute, Amsterdam, The Netherlands) and were maintained in Iscove’s Modified Dulbecco’s Media (IMDM, Thermofisher) supplemented with 10% FCS and P/S. DT-40 cells were grown in Roswell Park Memorial Institute (RPMI)-1640 media supplemented with 10% FCS, 1% chicken serum (Sigma) and P/S (Gibco) at 39.5°C, 20% O_2_ and 5% CO_2_. The KB2P3.4 cell line was established from a mammary tumor from K14cre;*Brca2*^F11/F11^;*p53*^F2-10/F2-10^ mice and described previously (Evers et al., 2008). The KB2P3.4R3 cell line was created by the stable introduction of an integrative bacterial artificial chromosome (iBAC), containing the full-length mouse *Brca2* gene, into the KB2P3.4 cell line (Evers et al., 2008). KB2P3.4 and KB2P3.4R3 cells were cultured in DMEM/F-12 medium, supplemented with 10% FCS, 50 units/mL penicillin, 50 μg/mL streptomycin, 5 μg/mL insulin (Sigma), 5 ng/mL epidermal growth factor (Life Technologies) and 5 ng/mL cholera toxin (Gentaur), at 37°C and hypoxic conditions (1% O_2_, 5% CO_2_).

### Knockdown and knockout cell line models

To generate RPE-1 and KBM-7 cell lines expressing doxycycline-inducible short-hairpin RNAs (shRNAs), DNA oligos were cloned into Tet-pLKO-puro (Addgene plasmid #21915) vector. Tet-pLKO-puro was a kind gift from Dmitri Wiederschain. shRNAs directed against luciferase (‘shLUC’, 5′-AAGAGCTGTTTCTGAGGAGCC-3′), BRCA2 (#1: 5′-GAAGAATGCAGGTTTAATA-3′ and #2: 5′-AACAACAATTACGAACCAAACTT-3′), BRCA1 (#1: 5′-CCCACCTAATTGTACTGAATT-3′ and #2: 5′-GAGTATGCAAACAGCTATAAT-3′), RAD51 (#1: 5′-CGGTCAGAGATCATACAGATT-3′ and #2: 5′-GCTGAAGCTATGTTCGCCATT-3′), MUS81 (#1 5′-GAGTTGGTACTGGATCACATT-3′ and #2 5′-CCTAATGGTCACCACTTCTTA-3′), SLX4 (#1 5′-ATTTCTGCTTCATTCACGTTT-3′ and #2 5′-CACCTGCAGACTCAAATGCCG-3′), and ERCC1 (#1 5′-CCAAGCCCTTATTCCGATCTA-3′ and #2 5′-CAAGAGAAGATCTGGCCTTAT-3′) were cloned into the Tet-pLKO-puro vector. Lentiviral particles were produced as described previously (Heijink et al., 2015). In brief, HEK293T packaging cells were transfected with 4□μg of indicated pLKO plasmid in combination with the packaging plasmids lenti-VSV-G and lenti-ΔVPR using a standard calcium phosphate protocol (van Vugt et al., 2010). Virus-containing supernatant was harvested at 48 and 72□h after transfection and filtered through a 0.45□μM syringe filter. Supernatants were used to infect target cells in medium with a final concentration of 4□μg/mL polybrene (Sigma Aldrich). RPE-1 cells harboring a *TP53* mutation were generated by introducing a single-guide RNA (sgRNA) targeting exon 4 of the *TP53* gene as described previously (Kok et al., 2020). To generate RAD52 and POLQ knockout cells, sgRNAs targeting exon 3 of RAD52 (AGAATACATAAGTAGCCGCA) and exon 1 of POLQ (GCCGGGCGGCGGGCTCAGCA) were cloned into the PX458 vector, which was a gift from Feng Zhang (Addgene plasmid # 48138). POLQ sgRNAs were a kind gift from Marcel Tijsterman (Leiden University Medical Centre, Leiden, the Netherlands). Plasmids were introduced in RPE-1 cells using Fugene HD transfection reagent and cells were selected based on GFP-expression or using 7 μg/mL puromycin (Sigma Aldrich) for 5 days.

### siRNA transfection

Cells were transfected with 40□nM siRNAs (Ambion Stealth RNAi, Thermofisher) targeting PARP1 (sequence 1: #HSS100243 and sequence 2: #HSS100244) or a scrambled (SCR) control sequence (sequence #12935300) with oligofectamine (Invitrogen), according to the manufacturer’s recommendations.

### Sister chromatid exchange assays

SCE assays were performed as described previously (Perry and Evans, 1975). RPE-1 cells were pre-treated with 0.1 μg/mL doxycycline (Sigma) for 48 h, followed by 48 h treatment with 10 μM BrdU. For BRCA2-deficient RPE-1 cells, BrdU treatment was increased to 64 h. Inhibitors were added for 48 h, simultaneously with BrdU treatment at the following concentrations: 0.5 μM olaparib (Axon Medchem), 16 μM veliparib (Axon Medchem), 7 nM talazoparib (Axon Medchem), 50 nM mitomycin C (Sigma), 5 μM cisplatin (Accord), 5 nM campthothecin (Sigma), 250 nM etoposide (Sigma), 20 μM BO2 (Axon Medchem). VE-821 (ATRi; Axon Medchem) was added simultaneously with colcemid for 3 h at a concentration of 1.0 μM. Alternatively, cells were treated with 2 Gy γ-irradiation 8-10 h prior to fixation using an IBL 637 Cesium137 γ-ray machine. Cells were collected in 10 μg/mL colcemid (Roche) for 4-6 h, fixed in 3:1 methanol:acetic acid solution and inflated in a hypotonic 0.075 M KCl solution. Metaphase spreads were made by dripping the cell suspensions onto microscope glasses from a height of ~ 30 cm. Slides were stained with 10 μg/mL bis-Benzimide H 33258 (Sigma) for 30 minutes, exposed to 245 nM UV light for 30 minutes, incubated in 2x SSC buffer (Sigma) at 60°C for 1 h, and stained in 5% Giemsa (Sigma) for 15 minutes. DT40 cells were treated with BrdU for 48 h, doxycycline for 24 h and 0.5 μM olaparib for 24 h. Alternatively, DT40 cells were treated with and BrdU and doxycycline as stated above, irradiated with 4 Gy and fixed a8 h later. KB2P3.4R3 cells were treated with BrdU for 32 h and KB2P3.4 for 40 h. Both KB2P3 cells lines were treated with 1 μM olaparib for 48 h.

### Immunofluorescence microscopy

RPE-1 cells were seeded on glass coverslips in 6-well plates and treated with doxycycline (1μg/ml) and olaparib (0.5 μM). Cells were then treated for 4 h with the CDK inhibitor RO-3306 (5μM). Upon washout of RO-3306, cells were incubated with EdU (20 μM) for 25 minutes. Cells were fixed using 2% formaldehyde in 0.1% Triton X-100 PBS for 10 minutes and subsequently permeabilized for 10 min in PBS with 0.5% Triton X-100. Staining was performed using primary antibodies against FANCD2 (Novusbio, Centennial, CO, USA; NB100-182, 1:200) and γH2AX Millipore, 05-636, 1:200). Cells were then incubated with corresponding Alexa-488 or Alexa-647-conjugated secondary antibodies and counterstained with DAPI (Sigma). For analysis of DNA damage response components, prophase and pro□metaphase cells were identified based on condensed chromatin conformation, and included for analysis. Images were acquired on a Leica DM6000B microscope using a 63× immersion objective (PL S-APO, numerical aperture: 1.30) with Las-af software (Leica, Wetzlar, Germany).

### Cell viability assays

RPE-1 cells were plated in 96-wells plates at a concentration of 800 cells per well. After 24 hours, cells were treated with indicated concentrations of olaparib, veliparip, or talazoparib (all from Axon Medchem) for 3 days. Methyl-thiazol tetrazolium (MTT, Sigma) was added to cells at a concentration of 5 mg/mL for 4 hours, after which culture medium was removed and formazan crystals were dissolved in DMSO. Absorbance values were determined using a Bio□Rad benchmark III Biorad microtiter spectrophotometer at a wavelength of 520 nm.

### Western blot analysis

Cells were lysed in Mammalian Protein Extraction Reagent (MPER, Thermo Scientific), supplemented with protease inhibitor and phosphatase inhibitor (Thermo Scientific). Protein concentrations were measured using a Bradford assay. Proteins were separated by SDS-PAGE gel electrophoresis, transferred to Polyvinylidene fluoride (PVDF, immobilon) membranes and blocked in 5% skimmed milk (Sigma) in TRIS-buffered saline (TBS) containing 0.05% Tween-20 (Sigma). Immunodetection was performed with antibodies directed against BRCA2 (Calbiochem, OP95, 1:1000), BRCA1 (Cell Signaling, 9010, 1:1000), RAD51 (GeneTex, gtx70230, 1:1000), PARP1 (Cell Signaling, 9532, 1:1000), RAD52 (Santa Cruz, sc-365341, 1:250), SLX4 (BTBD12; Novus Biologicals, NBP1-28680, 1:1000), MUS81 (Abcam, ab14387, 1:1000), ERCC1 (Cell Signaling, 3885, 1:1000), , HSP90 (Santa Cruz, sc-1055, 1:1000), and beta-Actin (MP Biomedicals, 69100 1:10000). Horseradish peroxidase (HRP)-conjugated secondary antibodies (DAKO) were used for visualization using chemiluminescence (Lumi-Light, Roche Diagnostics) on a Bio-Rad bioluminescence device, equipped with Quantity One/ChemiDoc XRS software (Bio-Rad).

### Strand-seq library preparation and sequencing

Strand-seq libraries were prepared as previously described (Falconer et al., 2012; Sanders et al., 2017), with a few modifications. Prior to sorting single cells, KB2P3.4 and KB2P4.4R3 were treated with 1μM olaparib and KBM-7 with 0.15 μM olaparib for 48 h. To incorporate BrdU during one cell cycle, BrdU (Invitrogen) was added to exponentially growing cell cultures at 40□μM final concentration. Timing of BrdU pulse was 16□h for KB2P3.4R3 and KBM-7 cells, and 20□h for KB2P3.4 cells. After BrdU pulse, cells were resuspended in nuclei isolation buffer (100□mM Tris-HCl pH 7.4, 150□mM NaCl, 1□mM CaCl2, 0.5□mM MgCl2, 0.1% NP-40, and 2% bovine serum albumin) supplemented with 10□μg/ml Hoechst 33258 (Life Technologies) and propidium iodide (Sigma Aldrich). Single nuclei were sorted into 5□μl Pro-Freeze-CDM NAO freeze medium (Lonza)□supplemented with□7.5% dimethyl sulfoxide, in 96-well skirted PCR plates (4Titude), based on propidium iodide and Hoechst fluorescence intensities using a FACSJazz cell sorter (BD Biosciences). For each experiment, 96 libraries were pooled and 250–450□bp-sized fragments were isolated and purified. DNA quality and concentrations were assessed on the Qubit 2.0 Fluorometer (Life Technologies) and using the High Sensitivity dsDNA kit (Agilent) on the Agilent 2100 Bio-Analyzer. Single-end 50 bp sequencing reads from the Strand-seq libraries were generated using the HiSeq 2500 or the NextSeq 500 sequencing platform (Illumina).

### Detection and mapping of breakpoints

Indexed bam files were aligned to mouse (GRCm38) or human genomes (GRCh38) using Bowtie254. Different R-based packages were used for the detection and mapping of breakpoints: Aneufinder2 was used for libraries with arbitrary copy number profiles (KB2P3.4 and KB2P3.4R3), while HapSCElocatoR (https://github.com/daewoooo/HapSCElocatoR) was used for libraries derived from the haploid cell line KBM-7. Aneufinder2 was used to locate and classify any type of breakpoint, not only template strand switches, using standard settings (Bakker et al., 2016). In short, copy numbers for both the Watson (negative) and Crick (positive) strand were called and breakpoints were defined as changes in copy number state. These breakpoints are then refined with read-resolution to make full use of the sequencing data. As Aneufinder2 also detects stable chromosomal rearrangements, clonal aberrations were defined as events that occurred at the exact same locations in >□25% of the libraries from one cell line. HapSCElocatoR is implemented in the R package fastseg (Klambauer et al., 2012), and uses circular binary segmentation to localize SCEs in haploid Strand-seq libraries as a change in read directionality from Crick to Watson or vice versa. Only non-duplicate reads with a mapping quality greater than or equal to 10 were analyzed. We considered only strand state changes with at least three directional reads on both sides of the putative SCE site as an SCE event. Single directional reads embedded within an extended region with the opposite directionality were considered as errors and their directionality was flipped. Computationally localized SCE or somatic copy number alteration (SCNA) events were further manually verified by visual inspection of chromosome ideograms (obtained from Aneufinder2 or BAIT; see Figure 1A and 5A respectively).

### Detection and analysis of SCE hotspots

HapSCElocatoR-generated ‘.bed’-files containing the locations of all mapped SCE events were uploaded to the UCSC Genome Browser and hotspots were identified as regions containing multiple overlapping SCEs. p-values were assigned to putative SCE hotspots using a custom R-script, based on capture–recapture statistics. Briefly, the genome was divided into bins of the same size as the putative hotspot and the chance of finding the observed number of SCEs in one bin was calculated based on the total number of SCEs detected in the cell line.

### Genomic analysis of SCE localization

A custom Perl script was used for the permutation model (https://github.com/Vityay/GenomePermute). For each of 1,000 permutations, a random number n was generated and all SCEs were shifted downstream by n bases on the same chromosome. To prevent small-scale local shifts, n was confined to be a random number between 2 and 50□Mb. When the resulted coordinate exceeded chromosome size, the size of chromosome was subtracted, so that the SCE is mapped to beginning part of the chromosome, as if the chromosome was circular. All annotated assembly gaps were excluded before our analysis, to prevent permuted SCE mapping to one of the gap regions. The number of SCEs overlapping with a feature of interest in each permutation was then determined, as well as the original SCE regions. All values were normalized to the median permutated value, in order to determine relative SCE enrichments over expected, randomized distributions and to allow for comparison of the different cell lines. Significance was determined based on the amounts of permutations that showed the same or exceeding overlap (enrichment) or the same or receding (depletion) overlap with a given genomic feature compared to overlap between the original SCEs and the same feature. Any experimental overlap that lies outside of the 95% confidence interval found in the permutations has a p-value below 0.05 and was deemed significant. Experimental overlaps lying outside of the permuted range were given a p-value below 0.001, as there was a <0.1% (1/1,000) chance of such an overlap occurring by chance.

Enrichment analyses for G4 motifs were performed using a 10□Kb SCE region size cutoff Putative G4 motifs were predicted using custom Perl script by matching genome sequence against following patterns: G3+N x G3+N x G3+N x G3+, where x could be the ranges of 1–3, 1–7, or 1– 12□bp. Enrichment analysis for coding genes, CFSs (HumCFS database), centromeres, and telomereswas performed used a 100□Kb size cutoff. Genome and gene annotations were obtained from Ensembl release 88 (GRCh38 assembly, http://www.ensembl.org). Gene bodies were defined as regions between transcription start sites and transcription end sites, gene promoters as 1□Kb regions upstream of transcription start sites.

### Flow cytometry

RPE-1 cells were treated with 20μM EdU for 0, 8, 12 or 24h, subsequently fixed in ice□cold ethanol (70%) for at least 16 h, and stained with primary antibody against phospho□histone□H3-Ser10 (Cell Signaling; 9701, 1:100) and Alexa□488□conjugated secondary antibodies (1:200). EdU Click-it reaction was performed with Alexa-647 azide according to the manufacturer’s instructions (InvitrogenTM). DNA was stained using propidium iodide following RNase treatment. At least 10,000 events per sample were analyzed on an LSR-II flow cytometer (Becton Dickinson, Franklin Lakes, NJ, USA). Data were analyzed using flowjo software (Becton Dickinson).

### Xenopus laevis egg extracts and biotin-oligonucleotide pull-downs

Cytostatic factor (mitotic) and low speed supernatant (LSS) extracts were prepared according to Murray and Blow respectively (Blow, 1993; Murray, 1991). Biotin-oligonucleotide pull-down MS was performed as previously described (Budzowska et al., 2015). In short, a biotinylated-oligo (5’-A*CGCTGCCGAATTCTACCAGTGCCTTGCTAGGACATCTTTGCCCACCTGCAGGTTCACCC-3‘, *=biotin) was annealed to its reverse complement at a concentration of 10 μM in 50 mM Tris pH8.0 buffer. The oligo-duplexes were diluted to 100nM, after which 10 μl oligo was coupled to 60 μl streptavidin-coupled magnetic beads (Dynabeads MyOne Streptavidin C1, Invitrogen) by incubation for 60 mins in wash buffer I (50 mM Tris pH 7.5, 150 mM NaCl, 1 mM EDTA pH8.0, 0.02% Tween-20). Excess oligo-duplexes were removed by three washes in IP buffer (ELB-sucrose buffer: 10mM HEPES-KOH ph7.7, 50mM KCl, 2.5 mM MgCl_2_, 250mM sucrose; 0.25 mg/mL BSA; 0.02% Tween-20), after which the oligo-beads mixture was suspended in 40 μl IP buffer. Mitotic and interphase extracts were thawed on ice from −80 °C and supplemented with 20x energy mix (20 mM ATP, 150 mM Creatine Phosphate, 20 mM MgCl_2_, 2.5 mM EGTA). For biotin-oligonucleotide pulldown 8 μl mitotic or interphase extract was incubated with 4 μl of oligo-beads mixture for 10 mins. Beads-extract mixture was washed two times with 400 ul of IP Buffer, two times with IP-buffer minus BSA, and lastly one time with ELB-sucrose buffer. After the final wash, beads were taken up in 50 μl denaturing buffer (8 M Urea, 100 mM Tris ph8.0) and snap frozen. Mass spectrometry of oligonucleotide-bound proteins was performed by on-bead digestion as previously described for plasmid pull-down MS (Larsen et al., 2019). Two biological replicate experiments were performed, each with three technical replicate measurements per sample.

### Mass spectrometry

Online chromatography of the extracted tryptic peptides was performed using an Ultimate 3000 HPLC system (Thermo Fisher Scientific) coupled online to a Q-Exactive-Plus mass spectrometer with a NanoFlex source (Thermo Fisher Scientific), equipped with a stainless-steel emitter. Tryptic digests were loaded onto a 5 mm × 300 μm internal diameter (i.d.) trapping micro column packed with PepMAP100, 5 μm particles (Dionex) in 0.1% formic acid at the flow rate of 20 μl/minute. After loading and washing for 3 min, trapped peptides were back-flush eluted onto a 50 cm × 75 μm i.d. nanocolumn, packed with Acclaim C18 PepMAP RSLC, 2 μm particles (Dionex). Eluents used were 100:0 H_2_O/acetonitrile (volume/volume (V/V)) with 0.1% formic acid (Eluent A) and 0:100 H_2_O/acetonitrile (v/v) with 0.1% formic acid (Eluent B). The following mobile phase gradient was delivered at the flow rate of 250 nl/min: 1–50% of solvent B in 90 min; 50–80% B in 1 min; 80% B during 9 min, and back to 1% B in 1 minutes and held at 1% A for 19 min which results in a total run time of 120 min. MS data were acquired using a data-dependent acquisition (DDA) top-10 method, dynamically choosing the most abundant not-yet-sequenced precursor ions from the survey scans (300–1650 Th) with a dynamic exclusion of 20 sec. Survey scans were acquired at a resolution of 70,000 at mass-to-charge (m/z) 200 with a maximum inject time of 50 milliseconds or AGC 3E6. DDA was performed via higher energy collisional dissociation fragmentation with a target value of 1×10E5 ions determined with predictive automatic gain control in centroid mode. Isolation of precursors was performed with a window of 1.6 m/z. Resolution for HCD spectra was set to 17,500 at m/z 200 with a maximum ion injection time of 50 milliseconds. Normalized collision energy was set at 28. The S-lens RF level was set at 60 and the capillary temperature was set at 250°C. Precursor ions with single, unassigned, or six and higher charge states were excluded from fragmentation selection.

### Electron microscopy analysis of DNA intermediates

Electron microscopy (EM) analysis was performed according to the standard protocol (Neelsen et al., 2014; Zellweger et al., 2015), with modifications. For DNA extraction, cells were lysed in lysis buffer and digested at 50 °C in the presence of Proteinase-K for 2 h. The DNA was purified using chloroform/isoamyl alcohol and precipitated in isopropanol and given 70% ethanol wash and resuspended in elution buffer. Isolated genomic DNA was digested with PvuII HF restriction enzyme for 4 to 5 h. DNA was washed with TE buffer and concentrated using Amicon size-exclusion column. The benzyldimethylalkylammonium chloride (BAC) method was used to spread the DNA on the water surface and then loaded on carbon-coated nickel grids and finally DNA was coated with platinum using high-vacuum evaporator MED 010 (Bal Tec). Microscopy was performed with a transmission electron microscope FEI Talos, with 4K by 4K cmos camera. Images were processed and analyzed using the MAPS software (FEI) and ImageJ software.

## Author contribution

A.M.H., P.L., and M.A.T.M.v.V. conceived the project. A.M.H., C.S, Y.K., A.A., R.H.d.B., M.E., E.W. conducted cell biological experiments. A.A., R.H.d.B. and P.K. coordinated and performed mass spec analyses using *Xenopus* egg extracts. D.S. coordinated sequencing analyses. D.P. and V.G. performed bioinformatics analysis. E.M.M. and A.R.C. conducted and analyzed ssDNA and EM analyses. A.M.H., C.S. and M.A.T.v.V. wrote the manuscript. All authors provided feedback on the manuscript.

## Data availability

Mass spec data is deposited at the PRIDE repository. StrandSeq data are deposited at the short-read archive (SRA) of NCBI.

## Acknowledgments

This work was supported by grants from the Netherlands Organization for Scientific Research (NWO-VIDI 917.13334 to M.A.T.M.v.V. and Gravitation program ‘CancerGenomiCs’ to P.K.), the European Research Council (ERC-Consolidator grant #681572 ‘TENSION’ to M.A.T.M.v.V.), ERIBA-UMCG funding to A.M.H., P.L. and M.A.T.M.v.V. We thank Jos Jonkers and Marcel Tijsterman for sharing reagents. We thank members of the Medical Oncology Department and ERIBA for feedback on the manuscript.

## Figure legends

**Figure S1: Olaparib-treatment induces somatic copy number alterations in *Brca2*^iBAC^ and *Brca2*^−/−^ cancer cells. (A)** Representative Strand-seq libraries of chromosome 6 in *Brca2*^−/−^ cells or *Brca2*^iBAC^ cells, treated with DMSO (top) or olaparib (bottom). Black arrowheads indicate copy number variations (CNVs), grey arrows indicate clonal aberrations.

**Figure S2: Olaparib-induced SCEs are dose-dependent and arise independently of canonical endonucleases. (A)** RPE1-*TP53*^−/−^ shBRCA2 cells were pretreated with doxycycline (dox) where indicated and subsequently treated with increasing doses of PARP inhibitors olaparib, veliparib and talazoparib for 3 days. Cell viability was measured by MTT conversion. **(B)** RPE1-*TP53*^−/−^ shBRCA2 cells were pretreated with doxycycline (dox) and treated with increasing doses of olaparib for 48 hours. Means and standard deviations of 30 mitoses per condition are plotted. **(C)** RPE1-*TP53*^−/−^ cells with indicated dox-inducible shRNAs were treated for 48 hours with doxycycline and immunoblotted for indicated proteins. **(D, E)** RPE1-*TP53*^−/−^ cells with indicated dox-inducible shRNAs were treated with doxycycline for 48 hours, and subsequently treated with olaparib for 48 hours (panel D) or 2 Gy irradiation (panel E). SCEs were determined by differential BrdU incorporation. Means and standard deviations of at least 30 mitoses per condition are plotted. **(F)** DT40 *RAD51*^−/−^ cells harboring a dox-repressed hRad51 transgene were treated with doxycycline for indicated time periods. Gaps and breaks are indicated with black arrows. Statistics in panels (B, D, E) were performed using unpaired two-tailed t-tests (ns: p > 0.05, *: p < 0.05, **: p < 0.01, ***: p < 0.001, ****: p < 0.0001).

**Figure S3: Mapping of olaparib-induced SCEs in KBM-7 cells. (A-D)** KBM-7 cells harboring doxycycline-inducible shLUC or shBRCA2 were pre-treated with doxycycline and subsequently treated with olaparib if indicated. SCEs (panel B), deletions (DELs; panel C), amplifications (AMPs; panel C) and copy number variations (CNVs; panel D) were scored using StrandSeq of 64 (shLUC/DMSO), 31 (shLUC/OLA), 50 (shBRCA2/DMSO) and 52 (shBRCA2/OLA) libraries per condition. **(E)** Representative scheme displaying SCE mapping based on the gap between the last Crick read and the first Watson read. **(F-I)** Mapping of SCEs to centromeres (panel F), telomeres (panel G), gene bodies (panel H) and G4 structures (panel I) were computed. Random permutations and observed SCEs are plotted. P values indicate deviation of the observed number of SCEs compared to the mean of all permutations.

**Figure S4: Characterization of RAD52 and POLQ-deficient cell lines. (A)** RPE1-*TP53*^−/−^ cells with dox-inducible shBRCA2 and CRISPR/Cas9-mediated RAD52 knockout were treated for 48 hours with doxycycline and immunoblotted for indicated proteins. **(B)** RPE1-*TP53*^−/−^sgRAD52 shBRCA2 cells were pretreated with doxycycline (dox) and subsequently treated with olaparib where indicated. SCEs were determined by differential BrdU incorporation. Means and standard deviation of 11-30 mitoses per condition are indicated. **(C)** *POLQ* mutation in RPE1-*TP53*^−/−^ shBRCA2 cells was assessed with TIDE and aligned to the reference genome, revealing a 1 bp insertion in exon 1. Statistics in panel (B) were performed using unpaired two-tailed t-tests (ns: p > 0.05, *: p < 0.05, **: p < 0.01, ***: p < 0.001, ****: p < 0.0001).

