## Supplementary figures and images for "Sister chromatid exchanges induced by perturbed replication are formed independently of homologous recombination factors"

### Supplemental Figure 1

#### Supplementary Figure 1

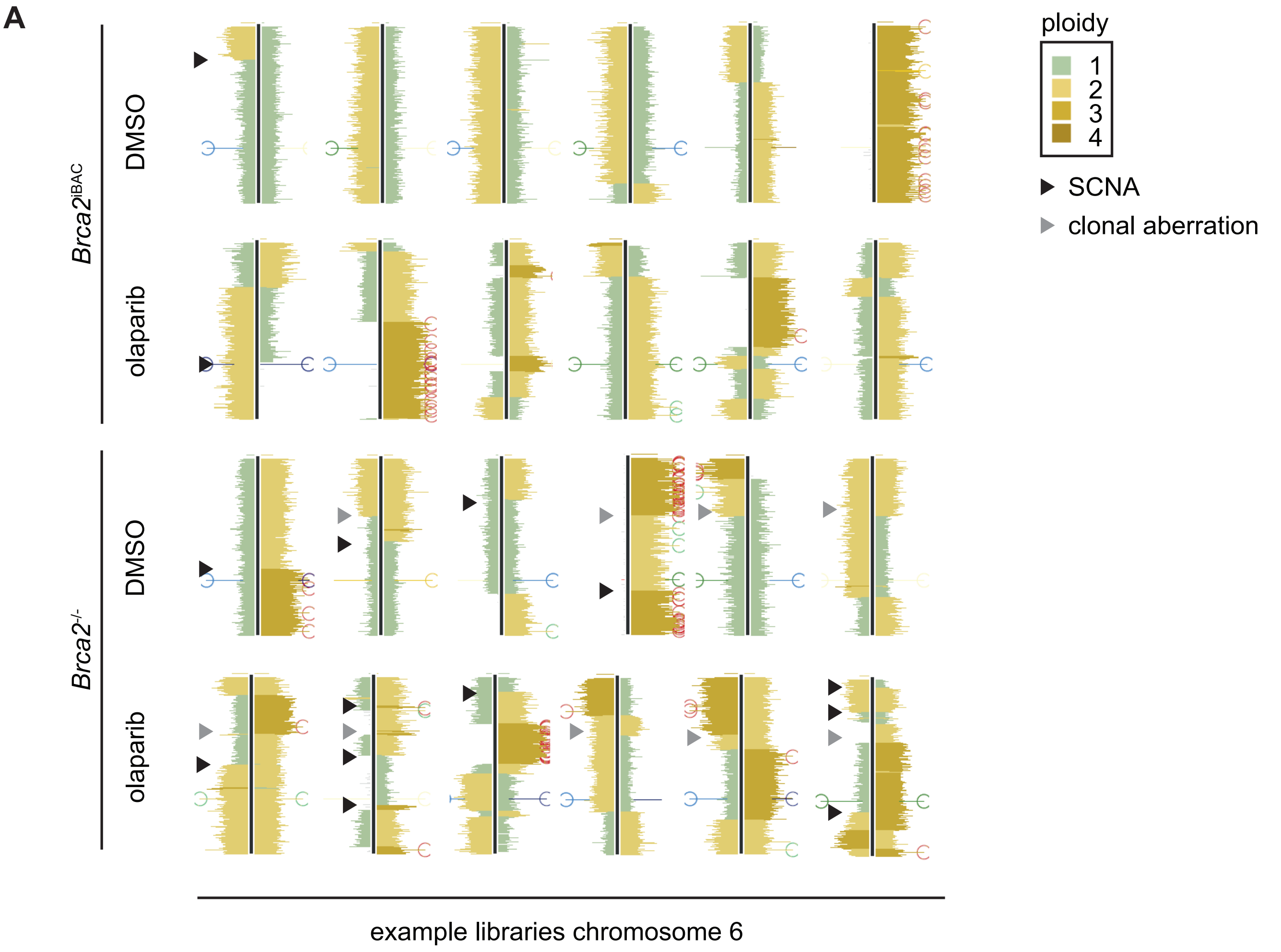

### Supplemental Figure 2

Supplementary Figure 2

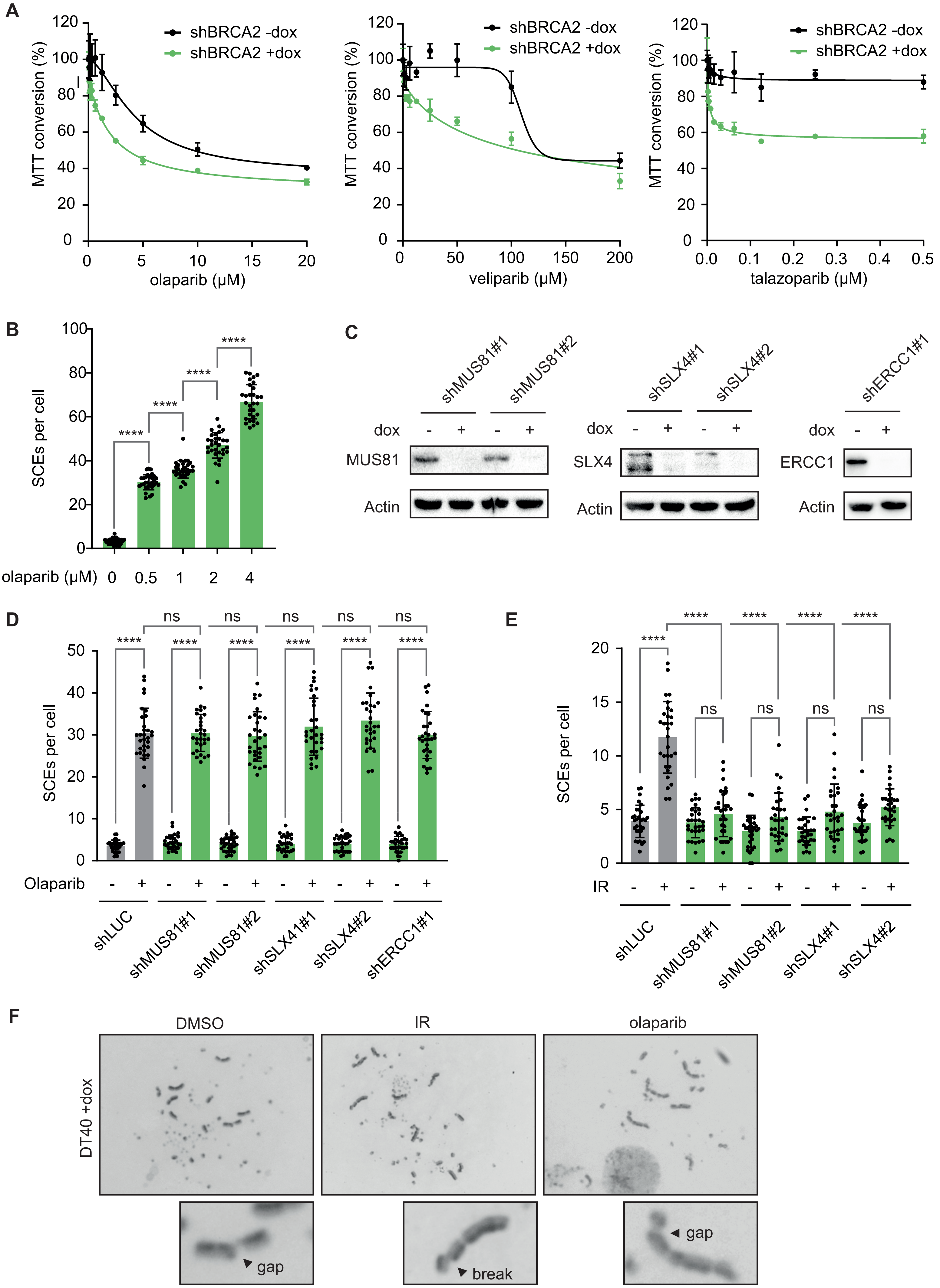

### Supplemental Figure 3

Supplementary Figure 3

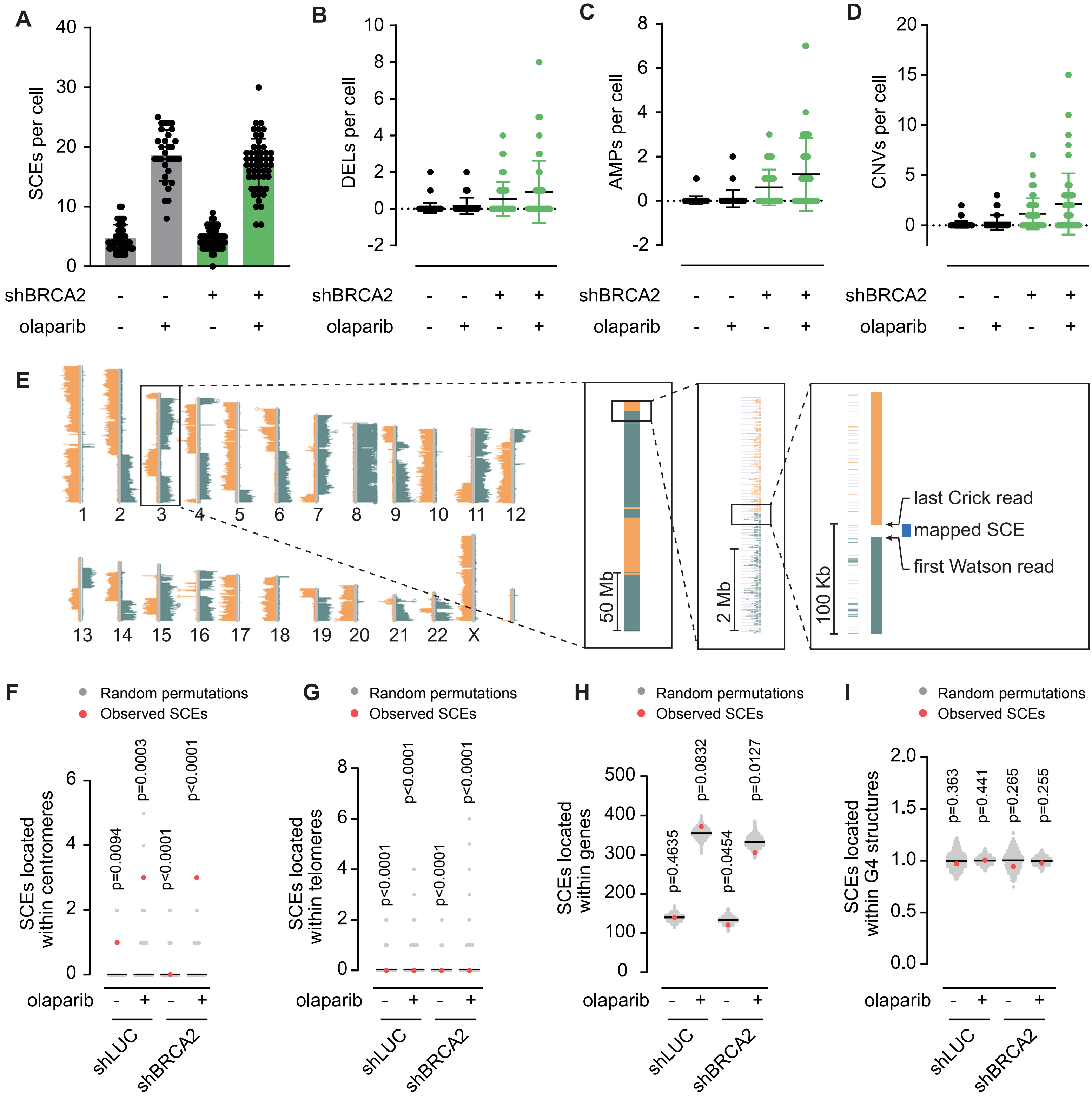

### Supplemental Figure 4

##### Supplementary figure 4

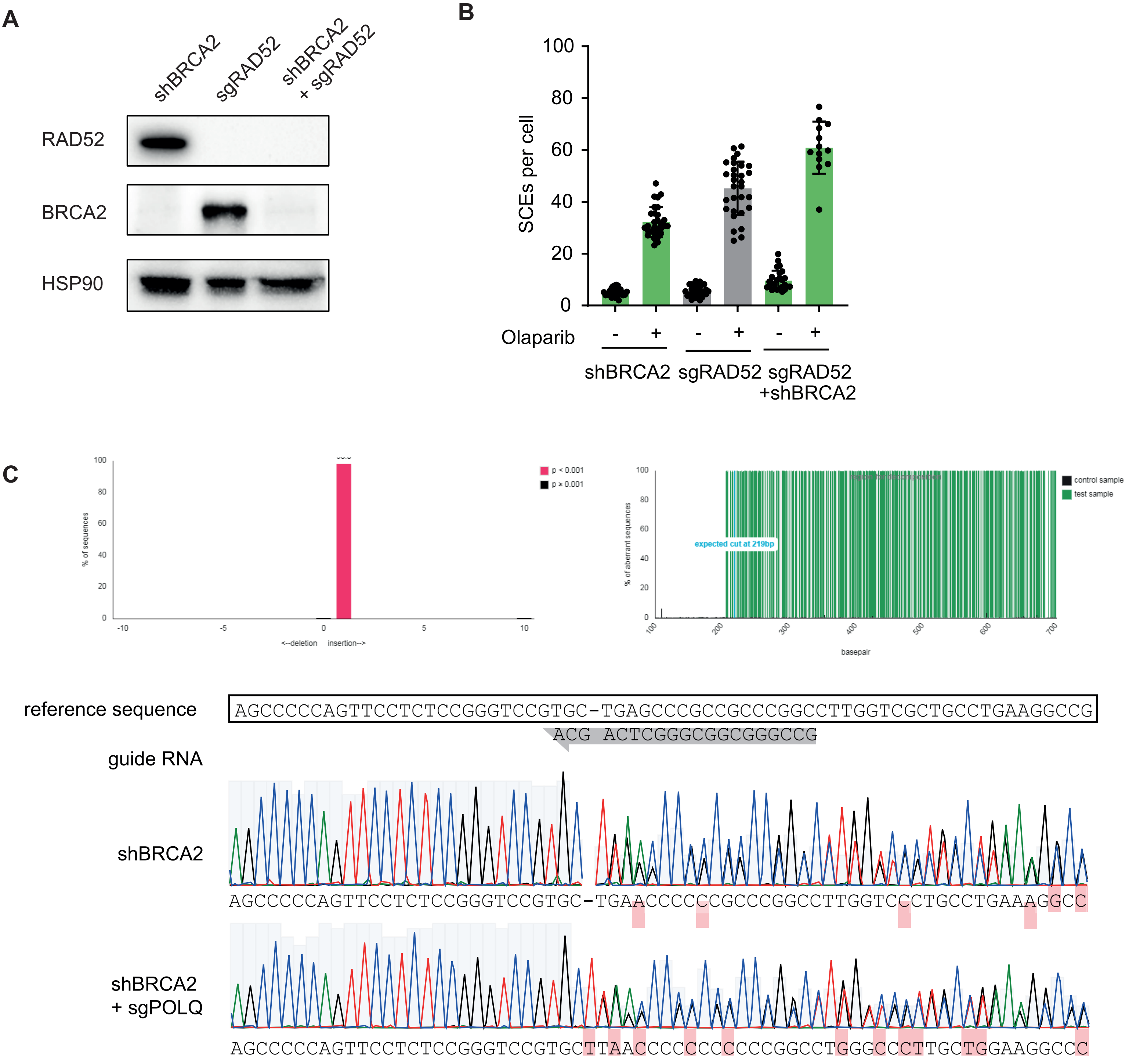
